# Cognitive Processing dissociation by mental effort manipulation in long demanding tasks

**DOI:** 10.1101/2020.04.25.060814

**Authors:** Marcus Vinicius Alves, Susanny Tassini, Felipe Aedo-Jury, Orlando F. A. Bueno

**Affiliations:** Department of Psychobiology – Universidade Federal de São Paulo, São Paulo, Brazil; Institute of Microscopic Anatomy and Neurobiology – Johannes Gutenberg University of Mainz, Mainz, Germany; German Resilience Center (DRZ), Mainz, Germany; Center of Mathematics, Computing and Cognition – Universidade Federal do ABC, Santo André/Brazil

**Keywords:** Pupillometry, Mental Effort, Processing Systems, Cognitive Control, Cognitive Resources.

## Abstract

Individuals uses cognitive resources to modulate performance in demanding tasks and a non-invasive and reliable way of measuring mental effort is pupillometry. This study aimed to test the mental effort related to different processing systems in long tasks with controlled and automatic demands. We conducted two experiments with healthy subjects: in Experiment 1 (n=15), using a metronome to ensure control on task pace, participants performed a serial number generation task (Counting; little to no effort tasks), a random number generation (RNG; effortful tasks), and no task (Unfilled interval; no effort at all). In experiment 2, (n=15) participants performed counting tasks with or without additional intermediary beeps produced by a metronome to assess the effect of a possible increase in effort demanded by the distractors. Experiment 1 showed differences between unfilled interval, counting and RNG. Experiment 2 showed that the intermediate beep made the counting tasks more demanding than the normal counting tasks. Notable in both experiments was the tendency of participants to demand mental effort at the beginning of the trial. These results indicate that previously effortless automatic tasks can become controlled, or at least more demanding, with a simple experimental manipulation. They also reveal that tasks that require mental effort over a long period will demand more than automatic ones, but even so the peak of this demand is in the initial trial period. Moreover, they reveal the high sensitivity of pupillometry for the measurement of mental effort employing different processing systems and cognitive resource modulation.

## 1. Introduction

Mental effort is the utilization of cognitive resources to process information, both in terms of speed and accuracy (Gopher and Donchin 1986; Kahneman 1973; Kool, 2018; Yeo and Neal 2008). Mental effort measurement is a valuable tool to explore hypotheses in experimental psychology and psychophysiology (Beatty 1982; Beatty and Lucero-Wagoner 2000).

Pupillometric studies provide evidence that the greater the effort undertaken to accomplish a task, the greater is pupil dilation (Eckstein et al. 2016; Hess and Polt 1964; Kahneman and Peavler 1969). Change in pupil diameter is considered to be an effective indicator of an individual’s mental activity (Hess and Polt 1964; Kahneman and Peavler 1969). Therefore, pupillary dilation provides a concomitant physiological index of brain activity for interpretation of psychological data (Kahneman and Peavler 1969; Mathôt 2018). Pupil dilation is commonly viewed as a very reliable and representative index of mental effort, with the peak of dilation in response to tasks usually happening approximately one second after the stimulus is shown to the participant (Wierda et al. 2012). Pupil dilation after baseline correction is usually used as pupillometric data, but pupillary dilation peak is also a robust index of mental effort, and in some cases even more reliable (Beatty and Lucero-Wagoner, 2000; Hershaw and Ettenhofer, 2018). Although not explicitly related to central cognitive processes, pupil dilation is empirically relevant and accurate for such measurements (Beatty and Lucero-Wagoner 2000; Kahneman 1973).

The relationship between pupil dilation and mental activity may be related to activity of noradrenergic neurons rising in the locus coeruleus (Jepma et al. 2011; Joshi et al. 2016; Murphy et al. 2014), implying the relationship between noradrenergic locus coeruleus neuronal activity in top-down attentional processes (Corbetta and Shulman, 2002), and consequently the LC-NE system – important for task engagement and performance (Aston-Jones and Cohen 2005) – may be related to different cognitive load and mental effort levels (Gratton, 2018; Oliva 2019). With this, the LC-NE system would be essential to “turn on” the cognitive system to ensure appropriate levels of activation for cognitive performance (Alnæs et al. 2014; Berridge and Waterhouse 2003; Kihara et al. 2015).

Some studies also have shown that LC activation is related with eye blink (Alnaes et al. 2014; Kihara et al. 2015). Eye blink rate (EBR) is a complementary measure of mental effort. Is theorized that eye blinks can reflect cognitive engagement as well, but only before a high demanding task begin (Siegle et al. 2008; Van Bochove et al. 2013). In view of that, EBR represent the preparation for doing a hard task. Pupil dilation is sometimes related with preparation for engagement in a demanding task (Moresi et al. 2008), and these two indices usually are related with effortful tasks (Fukuda, 1994; Fukuda et al. 2005; Ichikawa and Ohira 2004).

Mental effort is controlled by task demands, as more difficult tasks tend to require higher mental effort. Arousal caused by effort is modulated according to the accomplishment of the task, that is, cognitive feedback allows modulation of mental effort during tasks (Foroughi et al. 2017; Leppink & Pérez-Fuster, 2019; Sweller 1988; Yeo and Neal 2008). Tasks are assumed to require a fixed amount of cognitive resources to their realization, being these resources dependent of the cognitive load required by the difficulty of the task and by potential extra load related to distractions and poorly designed instructions. When these two added loads do not fill the entire capacity of an individual’s cognitive resources, the remainder resources can be used by the individual to better accomplish the task (Christie & Sharer, 2015). The achievement of this balance between demands to stabilize the resource allocation in face of distractors can be considered an important feature of this cognitive resource usage (Cools and D’Esposito, 2011; Zhang et al. 2015). Thus information processing is more flexible and adaptable.

Information is usually defined as being processed by individuals in two different systems, the first performing automatic processing and the second controlled processing (Schneider and Shiffrin 1977; Shiffrin and Schneider 1977). The automatic system processes information quickly, without awareness of individuals and requires low attentional resources, while the controlled system processes information slowly, with awareness, and requires high attentional resources (Hasher and Zacks 1984; Schneider and Shiffrin 1977).

It is theorized that the mental effort required to perform automated tasks is minimal and generally tasks related to automatism are easy or are tasks which have been assimilated by individuals after a considerable amount of training (McCabe et al. 2011). Automatic systems are related to implicit functions of the brain, necessary to perform faster and simpler tasks. Thus, the use of mental resources when performing automated tasks tends to be almost nonexistent or very low (Hasher and Zacks 1984; Schneider and Shiffrin 1977; Shiffrin and Schneider 1977).

New tasks tend to demand the use of greater cognitive resources, but over time, as they gradually become automatic, the resources required for successfully accomplishing the task become diminished. In the present study, we used simple number counting as the automatic tasks. Simple counting tasks have a low cognitive demand as counting is predefined and tend to quickly become automatic. More complex tasks require controlled processing and take longer than automatic tasks (Schneider and Shiffrin 1977; Shiffrin and Schneider 1977). The controlled system is associated with elaborate cognitive functions, allowing the emergence of the basic foundations for explicit memory, conscience and language.

Despite studies that relate these concepts, performance and cognitive load involved in tasks may not be directly linked (Isabella et al 2019; Mitra et al. 2017; van der Wel and van Steenbergen 2018). These indicate that ME in healthy subjects is not caused by an alteration of task engagement but is likely to be the consequence of a decrease in efficiency, or availability, of cognitive resources (Gergelyfi et al. 2015). Measuring pupil dilation can demonstrate when individuals have finally learned new tasks (Foroughi et al. 2017), studies on how tasks of different cognitive demands influence popular dilation are common, but little is studied on how task repetition and small manipulations can present themselves in pupil dilation. Also little is addressed how this dynamic is influenced by the duration and small manipulations in the tasks performed.

In the present study, we used the random number generation (RNG) task as the controlled system task because it is complex and requires a high level of attentional resources (Dirnberger and Jahanshashi 2010; Jahanshashi et al. 2006; Neuringer 1986), given that the RNG task demands control processing to randomize numbers (Towse and Neil, 1998). The RNG task may be applied with different levels of difficulty depending on the response rate required, reducing or augmenting the demand of mental resources of individuals (Dirnberger and Jahanshashi, 2010; Hamdan et al. 2004; Jahanshashi et al. 2006).

Recent studies have drawn attention to the possibility that processing systems do not exist in a duality, but in a continuum (Bugg et. al. 2008; Bugg and Crump 2012; Diede and Bugg 2017; Egner 2008). Using pupillometry, Diede and Bugg (2017) demonstrated an example of cognitive control without awareness. Their findings demonstrate that cognitive control may be fast, flexible, and operate outside of awareness, but not effortlessly. These findings support the idea of “automatic control” of the cognitive process. The term “automatic control” was coined to challenge the automatic/control dichotomy and is based on recent evidence that suggests that attentional control can occur automatically without conscious awareness (Jacoby et al. 2003).

In the present study, we tested the possibility of automatic processing being a subtle process that operates in a continuum. In addition, we aimed to test how the accomplishment of a task that does not demand a high level of cognitive resources is placed in this continuum. It was also one of our goals to see how the demand for mental effort in automatic (with and without manipulation) and controlled tasks would manifest over a longer period of time – given that eyetracking experiments tend to record only a few seconds per trial.

Another important point is the fact that although studies have shown that pupil dilation is related to awareness (Kang et al. 2015; Laeng et al. 2012) pupillometry is as yet unable to represent an index of processing dissociation that can separate automatic and controlled responses. Pupillometry is an important tool for differentiating decision-making processes, with pupil dilation being useful to analyze decision-making to engage in a task (Cavanagh et al. 2014; Massar et al. 2016), even in mice (Lee and Morgolis 2016) or in different processing system tasks (Bourisly 2015; van Der Mer et al. 2010). Response-conflict demands, when two incompatible responses are demanded, induce pupil dilation as cognitive control is needed to solve the problem (Cabestrero et al., 2009; Jainta and Baccino, 2010; Miyake et al., 2000). However, the consequences of response-conflict demand still are debatable, with various consequences still to be investigated, such as how individuals adjust their behavior to reduce resources costs (Paas and Ayres 2014; Rondeel et al. 2015).

While studies have demonstrated the validity of pupil dilation measurement to analyze mental effort as an important noninvasive technique for the measurement of mental effort (Just et al. 2003), including demonstrating different processing systems (Querino et al. 2015), there are few studies that seek to relate pupillometry and EBR data to dissociation of different processing systems. In addition, we intend to test the sensitivity of pupillometry for the measurement of increased cognitive demand by manipulating previously simple automated tasks.

In the present study, we verified how pupillometry is related to mental effort in tasks requiring top-down or automatic processes and to manipulate previously automated tasks in order to turn them into controlled tasks. We focused first on how pupillometry is related to different cognitive demands of the same task. The task we chose was the random generation of numbers because random generation is a goal-driven task that spins the activity of prefrontal/parietal areas involved with attentional top-down process (Jahanhashi et al. 2000), like the noradrenergic locus coeruleus neurotransmitter system. Second, we manipulated a known automatic task in order to see if it becomes a more (or less) cognitively demanding one by means of minor changes in it. This particular task consists in counting from 1 to 9 in the overtrained sequence of daily life.

Our hypotheses were that pupillometry demonstrates a considerable difference between automatic and controlled tasks, but with a small experimental manipulation, it would also be possible to perceive a variation in the automatic tasks that is an indicative of the higher mental effort, without these tasks necessarily being at the level of conscious awareness usually suggested for this to be a controlled task per se. In addition, we related the findings of this study with the capacity of individuals to allocate resources to perform tasks.

## 2. Materials and methods

### 2.1. Eyetracking Apparatus

To record participants pupil diameter we used the *T120*® *Tobii Eye-Tracker* device, developed by *Tobii Technology* (Danderyd, Sweden), with the following technical specifications: a 1280 × 1024 pixel, 30 × 22 × 30 inch flat screen monitor; a sampling rate of 120 Hz; an accuracy of 0.5 degrees; Binocular record; and freedom for head movement.

Given that the experimental focus was on pupillometric data and the eye tracker compensates for small movements of the head, the head movement of participants was not restricted. The eye tracker recorded the diameter of the right and left pupils simultaneously. The experiment was performed in a hermetically sealed room with artificial illumination – thus, there was no difference between the participants’ collection. The artificial lighting of the room was around 800 lux (a condition in which the pupil is at a baseline of about 4.5 cm)..

#### 2.1.1. Eye-Tracker Data Analysis

##### Pupil Dilation

Before the analysis, the data obtained by the eye tracker underwent a treatment in which any data from sessions failing to properly record the participant’s pupil for more than 15% of the recording time was excluded. At the beginning of the experiment, we recorded the participants’ baseline pupil diameters in order to subsequently perform the calculation to obtain the pupil dilation. During this baseline recording, the participants looked at the screen for three minutes and were asked to try not to think about anything or perform any mental effort.

Pupil dilation was averaged across both eyes, and this mean was used in our analysis. Pupil dilation in each condition was obtained by calculating the average pupil diameter in the experimental condition and subtracting the average baseline diameter. Pupil dilation outliers were removed using a trimmed mean calculus that excluded 10% of the extreme data (outliers and signal errors). In all participants, the trimmed mean was equal to the normal mean, ensuring a secure pupillary dilation mean.

We also gathered the pupil dilation peaks for further analysis. Pupil dilation peaks were reported on eye-tracker raw data, and also identified by visual inspection of the effort curves and checked in the correspondent time.

##### EBR

Eyeblinks were counted according to the following the criteria: 1) rapid, large changes (above 0.4 mm in 0.06 s) in pupil diameter; and 2) The eye trackers’ inability to register both eyes at the same time for 0.06 s. Any signal that did not match these criteria was considered an artifact and was removed from the data analysis. A brief explanation of this process can be found in the supplementary material.

### 2.2. Ethical Approval

The experiment was conducted in accordance with relevant regulations and institutional guidelines, and was approved by the local ethics committee of São Paulo Hospital, Universidade Federal de São Paulo. Informed consent was obtained from all individual participants included in the study.

## 3. Experiment 1

The first experiment was carried out with the intention of testing the participants’ pupil dilation while undertaking different processing systems tasks.

### 3.1. Tasks

#### Random number generation task

In this task, the participant was instructed to generate numbers between 1 and 9 randomly, avoiding the generation of predefined sequences (such as 3, 4, 5 or 9, 8, 7) (Schulz et al. 2012). This task requires the maintenance of several instructions and an understanding of the concept of randomness; integration of this information and its concomitant maintenance in working memory; adoption of strategies involving the selection of appropriate responses and inhibition of inappropriate responses; the monitoring of responses; and modification or alternation of the strategies employed, to maintain the randomness required by the task (Hamdan et al. 2004; Jahanshahi et al. 2006).

Three RNG conditions were used: Participants generated numbers at a rate of one number every 2 seconds, 1 second or 0.8 seconds (RNG-1/2s, RNG-1/1s and RNG-1/.8s, respectively). To verify if the participants were generating random numbers, their generations were assessed with the RNG calculator program to find the Evans’ index, a randomization rate index (Evans 1974; Towse and Neil 1998). As the Evans’ index depends on the number of performed generations, the minimum of 90 generations was established (in any condition) to calculate this index.

#### Serial Number Generation or Counting task

participants counted from one to nine repeatedly for three-minutes. The counting task has an ingrained logic and is easy for the participant to perform, with possible automation and a low use of cognitive resources. Four counting conditions were used. Participants counted one number every eight hundred milliseconds (C1/.8s), one number every second (C1/1s); one number every two seconds (C1/2s) and only odd numbers every second (C1/1s-odd).

#### Unfilled interval (UNF)

An interval in which the participants did not perform any task. This works as a control of participants’ cognitive demand when performing no task at all to compare with the task periods. The unfilled period is not the same as the baseline period. Both intervals are periods in which the participants did not perform task, however, baseline was recorded before any task, and the unfilled interval was recorded, mixed and randomized with the other tasks.

In all the tasks, participants generated numbers following a metronome (saying the numbers aloud with the beep of the metronome), with the exception of the Unfilled interval.

The calculation of the Effect Size was performed with the open software Effect Size Generator – Professional Edition 4.1. The index to represent Effect Size was Cohen’s d, which may indicate a small (0.2-0.3), medium (about 0.5) or large (from 0.8) effect (Cohen 1994; Steiger 2004). For paired and repeated measures data, the d index was corrected from the difference of the means and original standard deviations (Dunlop et al. 1996).

### 3.2. Participants

The test group comprised 16 volunteers of both sexes, but two participants were excluded as their pupillometric data did not meet our data criteria as described above, leaving 15 participants (mean age 24.38 (± 3.50); and mean education years 14 (± 2.55), 11 being women (73.33%).

Exclusion criteria were: drug use history or treatment with drugs acting on the central nervous system, a history of psychiatric or neurological disorders, use of ophthalmic drugs or present ophtalmoparesis or ophthalmoplegia. A structured health screening questionnaire was completed by the participants to ensure that they met the eligibility criteria.

### 3.3. Procedure

Each experimental session was conducted individually and lasted about 40 minutes. Participants were seated comfortably in front of the eye-tracker’s screen at a distance of approximately 65cm. An individual nine-point calibration was performed. The screen remained black during the experiment, with a small fixation point (a white cross) in the center which the participants were asked to focus on when performing the tasks.

After recording the participant’s baseline pupil and explaining the tasks to be performed, the participant performed each task in a pseudo randomized order – different for each participants – for three minutes each, with a one-minute rest interval between tasks. During this interval, the participant was instructed to continue to focus on the screen. In the unfilled interval the participants did not perform any task, and they were instructed to try not to think about anything and just relax while looking at the screen.

### 3.4. Results

A repeated measures ANOVA to analyze pupil dilation in the various conditions revealed a statistical difference [F (7; 98) = 16.733, *p* <.001], and the Bonferroni’s *post-hoc* test showed a difference between the RNG conditions and all other conditions (Counting tasks and unfilled interval) [*p* < .05; d > 1.0]; and the unfilled interval and all the other conditions [p<.05; d > .6], except for the condition C1/1s [p=.424] (Fig 1).

**Fig 1:**
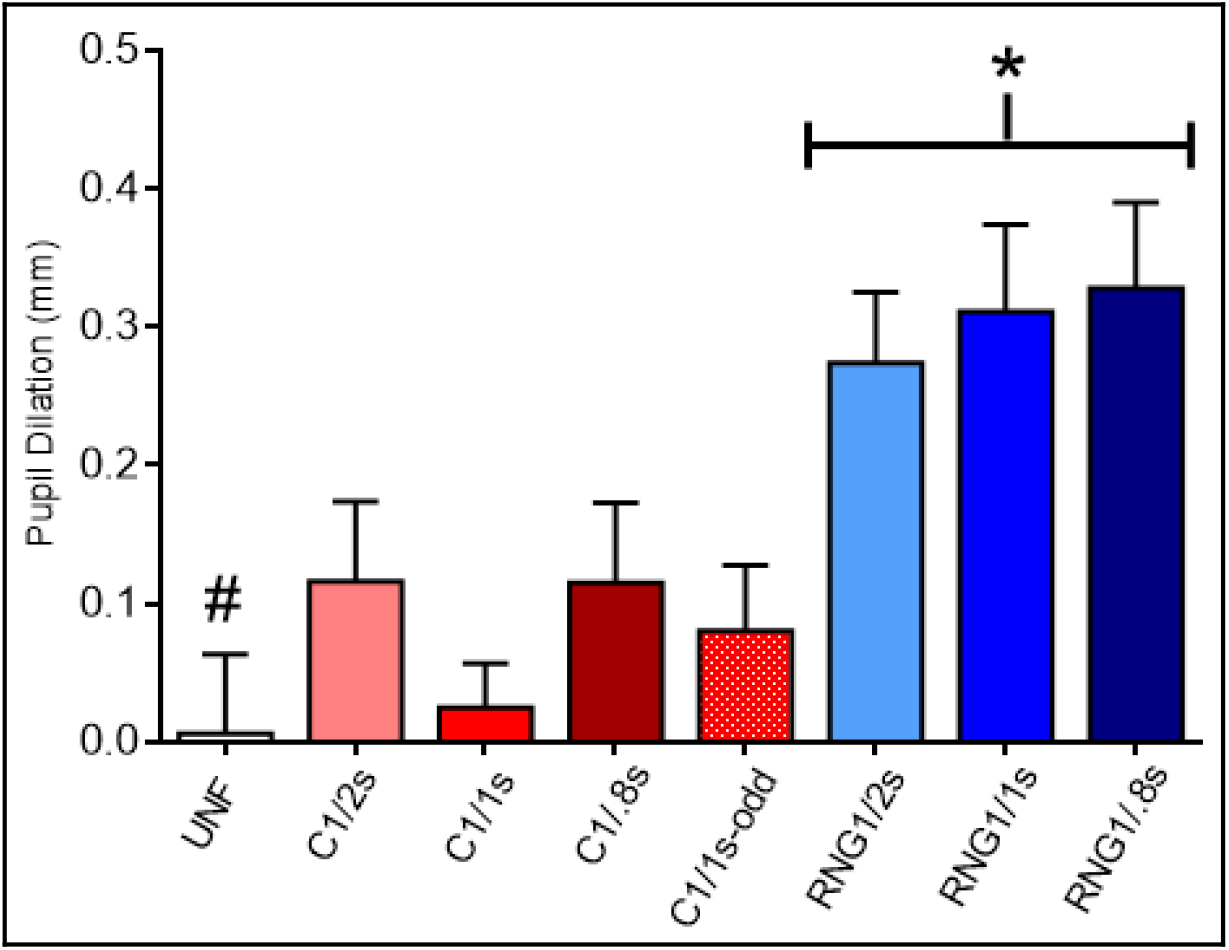
Pupil dilation in millimeters per condition showing differences between the unfilled interval, automatic (except C1/1s), and controlled tasks. Conditions are UNF (unfilled interval), counting task with pace of one generation per two seconds, one second, and 0.8 seconds (C1/2s, C1/1s, and C1/.8s, respectively), counting task of one odd number per second (C1/1s-odd), and random number generation tasks with one generation per two, one, and 0.8 seconds (RNG1/2s, RNG1/1s, and RNG1/.8s, respectively). * Represents statistical difference (p<.05). We used # to represent statistical difference (p<.05) to between the unfilled interval (UNF) and all other conditions, except for C1/1s. The bars represent standard errors.

In the repeated measures ANOVA for the peaks of pupil dilation, we found a statistical difference between conditions [F (7; 98) = 8.013, *p* <.001]; Bonferroni’s *post-hoc* test showed differences between RNG conditions and the others [*p* <.001; d > 1.0], except C1/1s-odd [p=.45 to RNG1/2s; p=.24 to RNG1/1s; and p=.62 to RNG1/.08s] and C-1/2s [p= 1.0 to RNG1/2s; p= 1.0 to RNG1/1s; and p=1.0 to RNG1/.08s]. Dilation peaks were found at the beginning of the trials, when the participants are more directed to the accomplishment of the tasks – more motivated. Moreover, the analysis of EBR did not show any difference between conditions (Table 1).

**Table 1:** Means (±SD) of Pupil dilation peak in millimeters and Eye Blink Rates (EBR) per conditions in Experiment 1

|  | UNF | C-1/2s | C-1/1s | C-1/.8s | C1/1s-odd | RNG-1/2s | RNG-1/1s | RNG-1/.8s | p |
| --- | --- | --- | --- | --- | --- | --- | --- | --- | --- |
| Pupil peak | .02<br>(.30) | .43<br>(.30) | .18<br>(.23) | .26<br>(.32) | .34 (.31) | .41<br>(.20) | .41 (.27) | .33<br>(.32) | **<.05** |
| EBR | 20.1<br>(11.0) | 19.1<br>(9.8) | 19.6<br>(10.6) | 16.8<br>(12.2) | 22.1<br>(8.7) | 18.5<br>(9.9) | 20.1<br>(9.6) | 18.9<br>(10.9) | .78 |

A Pearson correlation was performed to determine the relationship between the randomization index (Evans’ index) and pupil dilation in the RNG conditions. Despite the strong correlation between RNG conditions (RNG-1/2s and RNG-1/1s [r = .889, p <.001]; RNG-1/2s and RNG-1/.8s [r = .915, p <.001]; and RNG-1/1s and RNG-1/.8s [r = .915, p <.001]), we did not find any correlation with the random number generation index (RNG-1/2s [r = .087, p = .74]; RNG-1/1s [r = .325, p = .20]; RNG-1/.8s [r =.310, p =.23]).

An ANOVA was performed to verify the difference between conditions per time (1^st^ x 2^nd^ x 3^rd^ minute) for both pupil dilation and for EBR. Here we verified the difference within the conditions during the time performing the tasks. Regarding pupil dilation, we found statistical difference in all conditions between the three minutes, being the first minute the peak of pupil dilation and the third minute the minimum dilation, except for the second and third minutes in UNF [F(2, 30)=17.806, p=.158] (Fig2). Regarding EBR, there was no statistical difference regarding the time during the experiment [p >.05] (Fig3).

**Fig 2:**
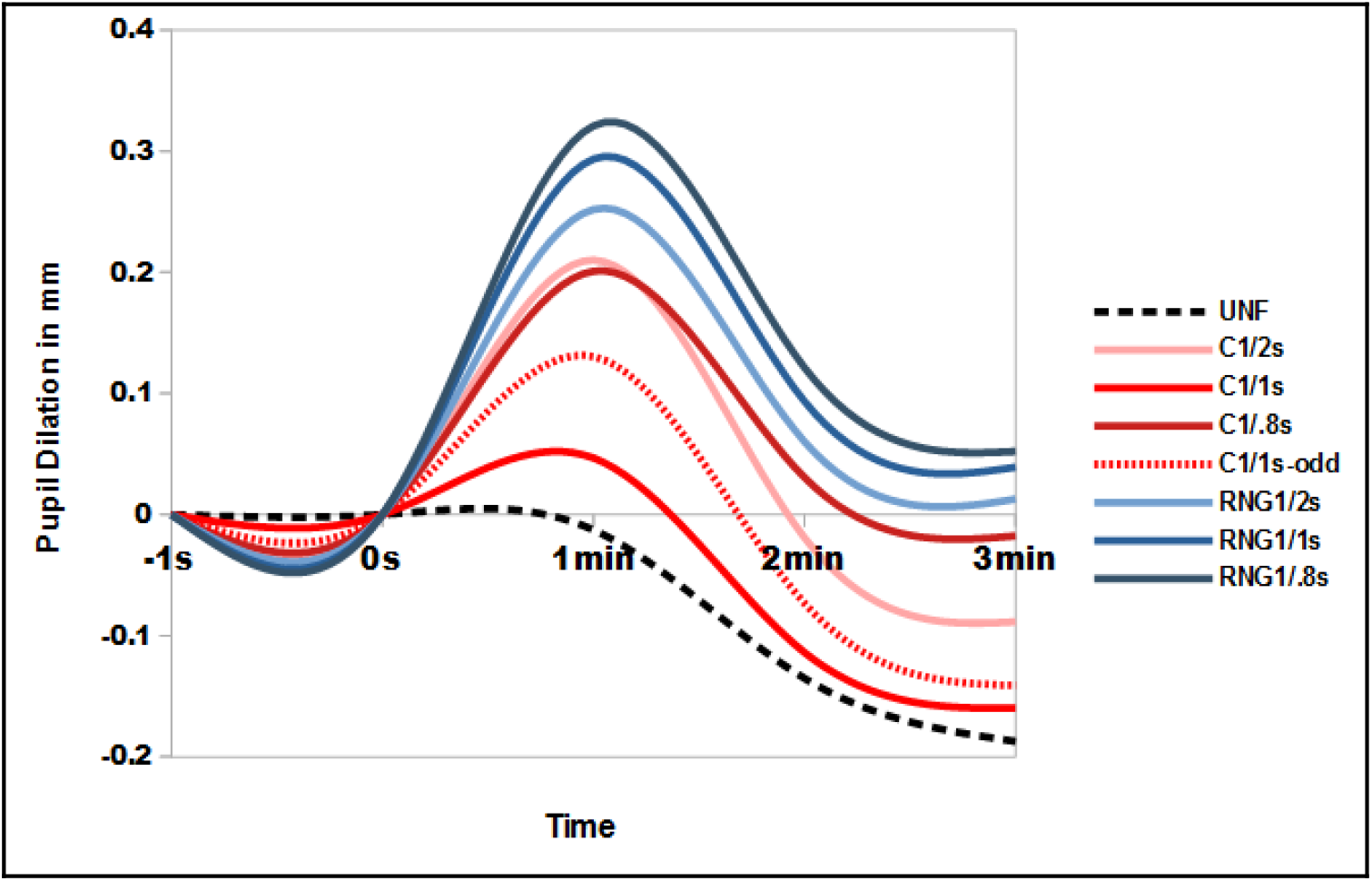
Pupil dilation in millimeters per condition during the 3-minutes trial. Lines represent the conditions. We verified the difference within the conditions during the time performing tasks (1^st^ to 3^rd^ minute). The analysis performed here was within the conditions and not between conditions. There was a statistical difference between the means over time in almost all conditions, with the first minute being the peak of pupillary dilation and the last minute always being the minimum of pupillary dilation. The only condition that did not have this result was the condition UNF, in this case there was difference only between the first minute and the other two minutes, the second and third minutes not requiring different mental effort levels in relation to pupillometry.

**Fig 3:**
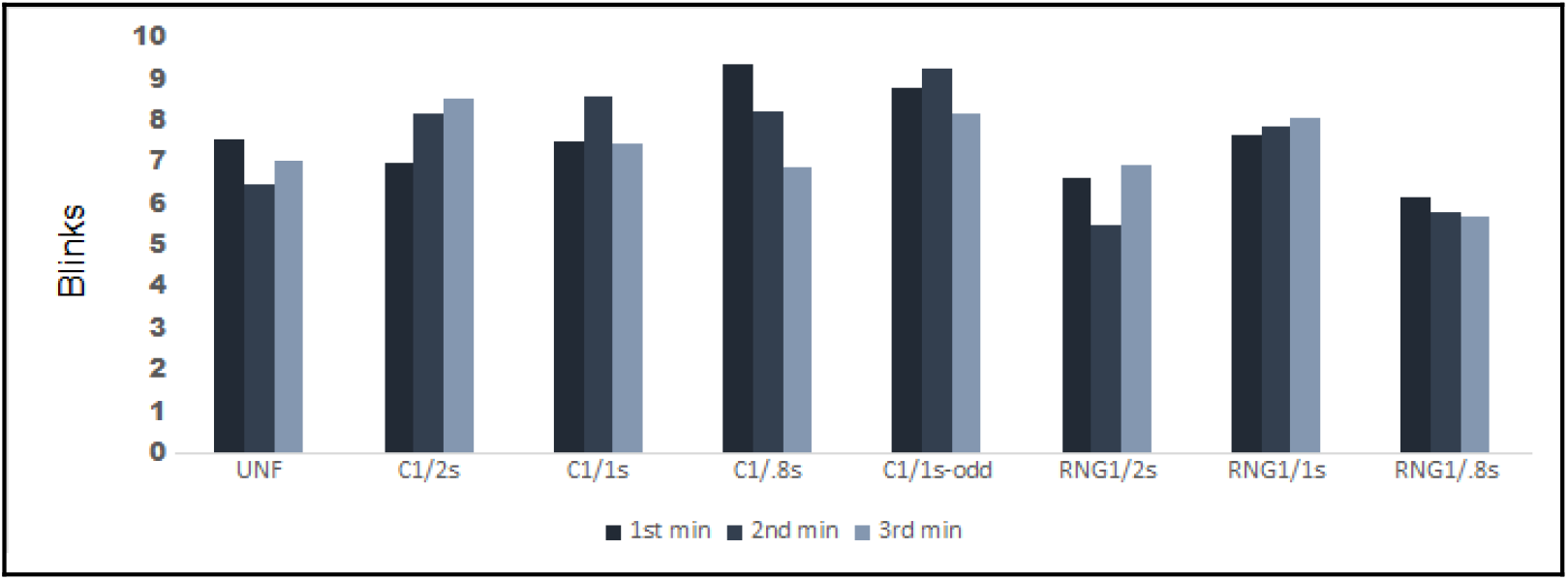
Eye blink rate (EBR) per conditions during the 3-minutes experiment trial. Bars represent the means of ERB. There was no statistical difference between conditions during the trials.

### 3.5. Discussion

These results indicated differences in pupil dilation between the unfilled interval, the counting tasks (associated with automatic processing) and the RNG tasks (associated with controlled processing). Considering pupil dilation as a measure of cognitive demand, the RNG tasks were all equally demanding, and were more demanding (higher pupil dilation) than the counting tasks and the unfilled interval. Therefore, effortless and effortful processes can be clearly differentiated by the use of pupillometry, as demonstrated in previous literature (Beatty and Lucero-Wagoner 2000; Kahneman 1973). But even automatic tasks (counting 1 to 9) may demand more resources than an unfilled interval. The only exception was found in condition C1/1s (counting one number per second), which was similar to the infilled period. This can be explained by the easiness of this particular task that makes it to require so little mental effort equivalent to an unfilled interval (Posner and Fan 2008). The counting task seems to have an optimal rate, perhaps the same as counting that people carry out on a daily basis. Of course, more evidence in needed to certify this assertion. Nonetheless, the counting task of one number per second can be used as a control condition to the unfilled period, avoiding participants’ reverberation of information or even mind-wandering in experiments in experiments involving an unfilled interval.

The results for the pupil dilation peaks during our tests corroborate these findings, differentiating again tasks with low cognitive demand and those with high cognitive demands. However, the counting tasks peaks were similar to the unfilled interval peaks, except for counting of odd numbers, one number per second (C1/1s-odd condition). The peak signals the start of attentional demand and usually occurs at the beginning of the tasks (Siegle et al. 2008). This result implies that the counting tasks – probably because of their already ingrained logic – do not oblige the participant to adjust their behavior at the beginning of the task, whereas the RNG tasks do demand this. This idea also explains the different result of the C1/1s-odd condition, considering that, although it requires low cognitive reallocation and preparation, it is not as simple as counting from 1 to 9 as the participant must inhibit the urge to count pair numbers leading to a greater likelihood of a pupil dilation peak.

Another point is that the pupil dilation peaks occur at the beginning of the experiment while the participant is still adapting to the task and accompanying the metronome. As has already been shown, a short time after the start of the task is the time when people make adjustments to produce a better performance (Wierda et al. 2012). It is possible that during the accomplishment of the tasks the participants are not only seeking to perform the tasks with more effectiveness, but are also adjusting the use of cognitive resources from the feedback from this accomplishment and, therefore, accomplishing the tasks using the least of cognitive resources possible (Debue and van de Leemput 2014; Yeo and Neal 2008).

The lack of differences between the EBR among the conditions does not necessarily imply that the use of cognitive resources was similar. One possible explanation for this phenomenon is that blinking is actually related to cognitive engagement (Jongkees and Colzato 2016), and thus, after the task is started the participants stabilize the amount of blinking during the three minutes of the task. Thus, there were not enough blinks for a statistical effect. Another possibility, which would go against the literature on blinking, is that this effect is not so susceptible to cognitive demands, at least to the demands of the tasks of this study. Since blinking tends to be considered representative of a number of factors, it is still far from being well defined theoretically (Debue and van de Leemput 2014).

Given that the pace of random number generation was not significantly different, it seems that the rate of generation is not the main indicator for the difference in mental effort. That is, the generation itself is cognitively demanding not the efficacy or quantity of the generations.

## 4. Experiment 2

In this experiment, an experimental manipulation was introduced in the counting tasks in order to add an extra demanding requirement. In the regular task in experiment 1, the subject was asked to count repeatedly from 1 to 9 paced by a metronome beep. We reasoned that another beep between the pacing beeps would demand more attentional resources because this intermediate beep is indeed a distracting stimulus that must inhibit the participant’s ability to keep the right rate.

This second experiment thus had two main goals; first, to replicate the aforementioned results of the counting tasks, and second, to verify if counting tasks with intermediate beeps would become more demanding, thus, requiring increased cognitive control.

### 4.1. Tasks

In this experiment, the participants performed the counting tasks only, but under two different conditions: Continuous counting (without intermediate metronome beeps) as in the first experiment; and intermittent counting (with an intermediate beep between target counts).

The continuous counting rates were one count per second (cC1/1s), one count every two seconds (cC1/2s), and one every three seconds (cC1/3s). The intermittent counting with intermediate beeps was at the same rate as the continuous counting, but the beeps turned the task into a go/no-go task (count/no-count/count): one count per second (iC1/1s), with beeps every 0.5 second; one count every two seconds (iC1/2s) with beeps every second; and one count every three seconds (iC1/3s-A), with beeps every 1.5 seconds. Moreover, we added one condition with beeps between the counting to verify if the pupil dilation effects that we observed were the result of a pace effect or an inhibitory effect: iC1/3s-B, in which the participants counted every three seconds, with one beep per second (count/no-count/no-count/count).

### 4.2. Participants

Fifteen volunteers of both sexes (mean age 22.87 (± 4.79); and mean education years 14.07 (± 1.91)), of whom 11 were women (73.33%). The same criteria as in the first experiment were considered for exclusion.

### 4.3. Procedure

The procedures in this second experiment were the same as in the first one.

### 4.4. Results

A repeated measures ANOVA for pupil dilation showed a statistical difference between conditions [F (6; 84) = 3.817, *p* <.05]. Bonferroni’s *post-hoc* test showed differences only between the conditions iC1/3s-B, iC1/2s and iC1/1s and the other conditions [p<.05; d = .5], with the exception of the condition iC1/3-B, that had a small effect when compared with condition iC1/3s-A (d = .3)]; the condition iC1/3s-A was not different from the continuous counting conditions (Fig 4).

**Fig 4:**
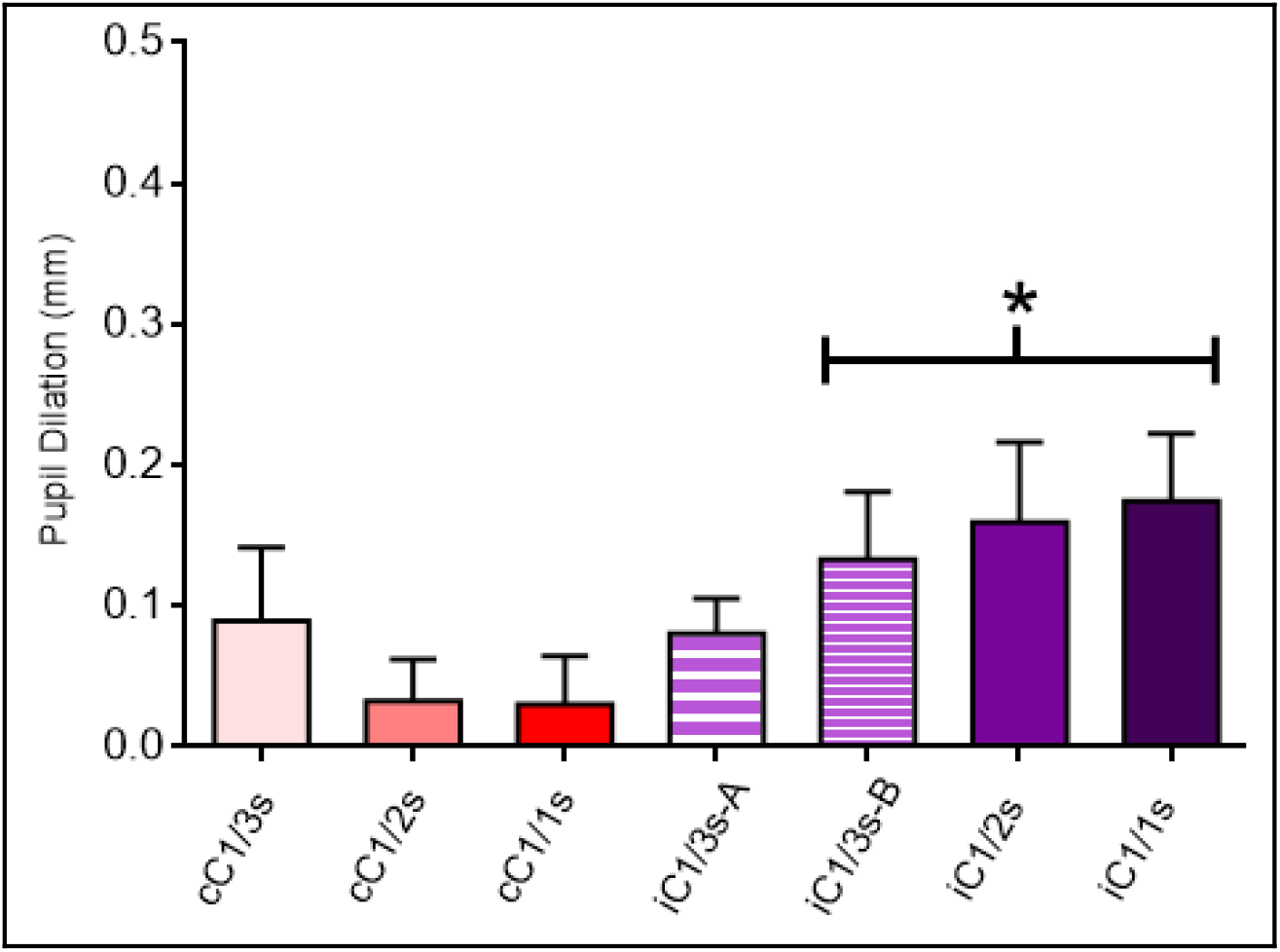
Pupil dilation in millimeters per condition showing differences between the continuous counting tasks – automatic tasks – and the intermittent counting tasks (except iC1/3s-A) – manipulated tasks. Conditions are counting tasks with pace of one generation per three, two, and one second (cC1/3s, cC1/2s, and cC1/1s, respectively), and counting tasks with one generation per three, two, and one second, with one beep (iC1/3s-A, iC1/2s, and iC1/1s respectively) or two (iC1/3s-B) as a distractor. * Represents statistical difference (p<.05). The bars represent standard errors.

In the repeated measures ANOVA for the peaks of pupil dilation, we found a statistical difference [F (6; 72) = 4.720, p < .001]; Bonferroni’s post-hoc test only showed differences between both the one count per second tasks: cC1/1s and iC1/1s [p < .001; d > 1]. Again, the repeated measures ANOVA for EBR showed no difference between conditions.

**Table 2:**
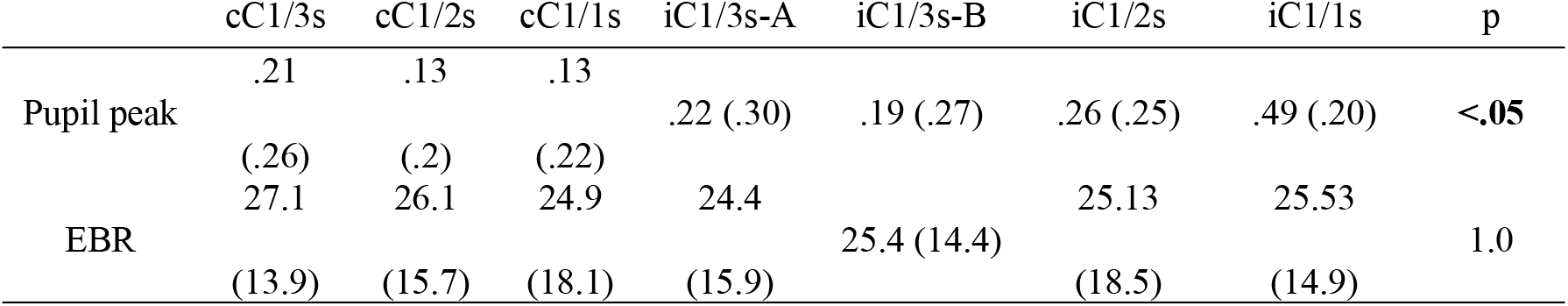
Means (±SD) of Pupil dilation peak in millimeters and Eye Blink Rates (EBR) per conditions in Experiment 2

|  | cC1/3s | cC1/2s | cC1/1s | iC1/3s-A | iC1/3s-B | iC1/2s | iC1/1s | p |
| --- | --- | --- | --- | --- | --- | --- | --- | --- |
| Pupil peak | .21<br>(.26) | .13<br>(.2) | .13<br>(.22) | .22 (.30) | .19 (.27) | .26 (.25) | .49 (.20) | **<.05** |
| EBR | 27.1<br>(13.9) | 26.1<br>(15.7) | 24.9<br>(18.1) | 24.4<br>(15.9) | 25.4 (14.4) | 25.13<br>(18.5) | 25.53<br>(14.9) | 1.0 |

In this experiment we also did an ANOVA that was performed to verify the difference between the conditions per time (1^st^ x 2^nd^ x 3^rd^ minutes) both to pupil dilation and EBR. Here we verify the difference within the conditions during the time performing the tasks. Regarding pupil dilation, there was a difference in all conditions from the first to the second and from the second to the third minute, except in iC1/3-A [F(2, 26)=78.870, p=.905] (Fig5), in this condition the second and third minutes weren’t different.

**Fig 5:**
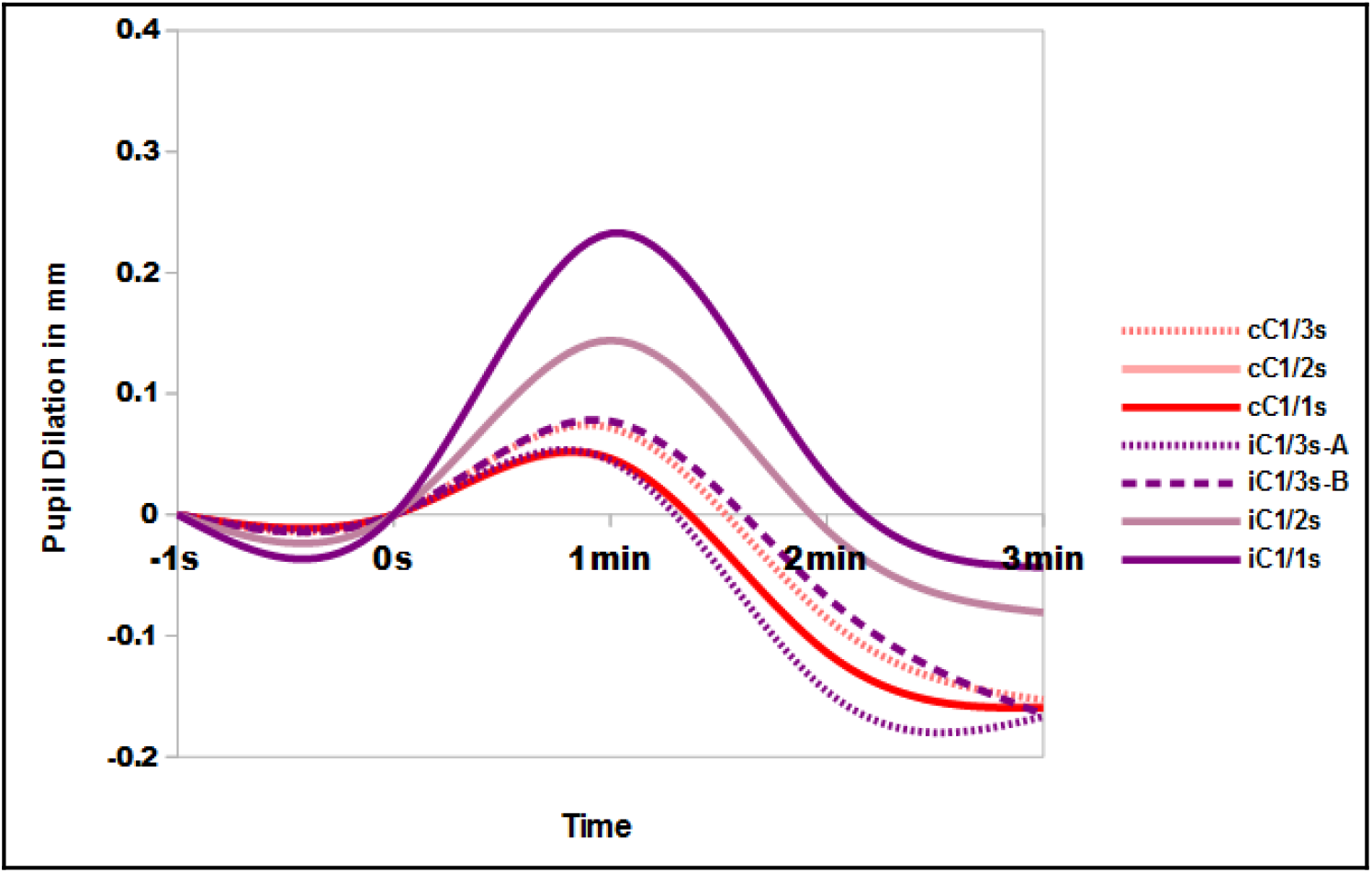
Pupil dilation in millimeters per condition during the 3-minutes trial. Lines represent the conditions. We verified difference within the conditions and not between conditions. There was a statistical difference between the means over time in almost all conditions, with the first minute being the peak of pupillary dilation and the last minute always being the minimum of pupillary dilation. The condition that did not have this result was iC1/3s-A, since the second and third minutes were not different.

Regarding EBR we only found statistical difference between the first minute and the later minutes in condition iC1/3-A, since there was no difference between the 2^nd^ and 3^rd^ minutes [F(2, 26)=5.781, p=.158] (Fig6). In all the other conditions there was no statistical difference on time during performing tasks and EBR.

**Fig 6:**
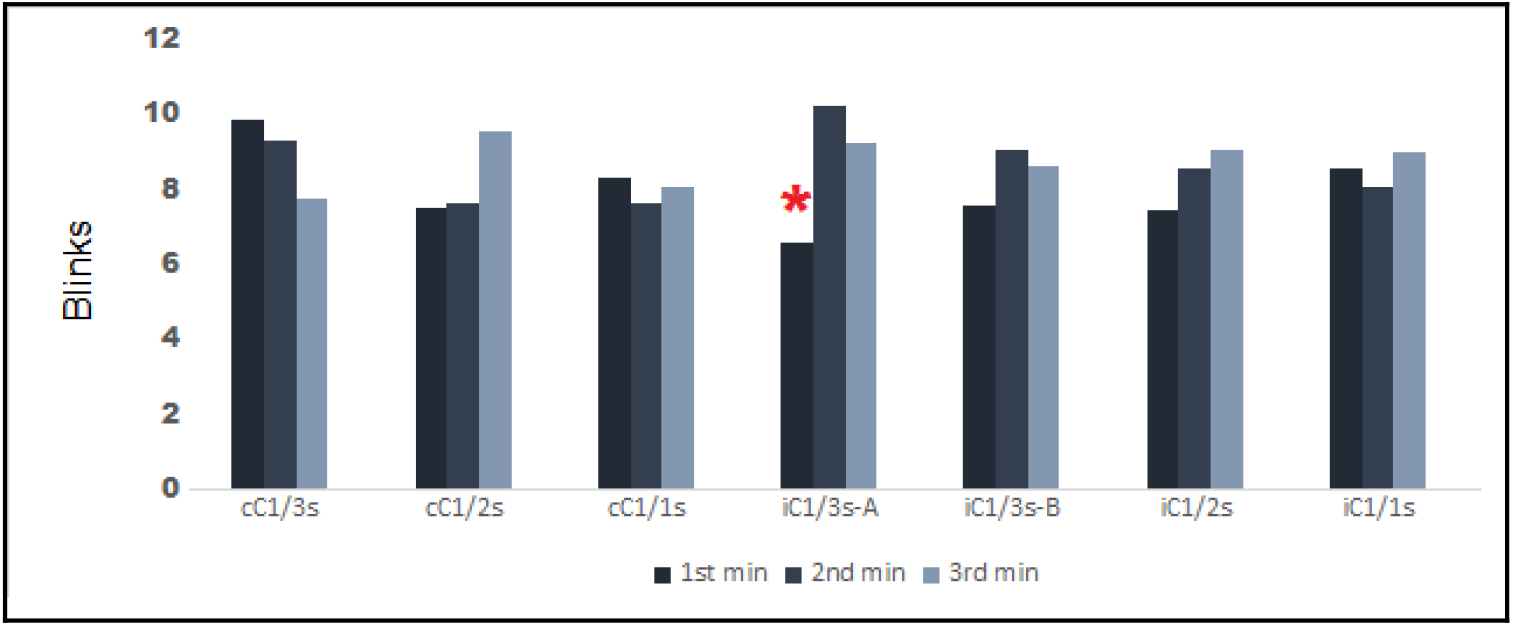
Eye Blink Rate (EBR) mean per conditions during the 3-minutes experiment trial. Bars represent means of ERB. The * represents a difference (p<.05) between the EBR mean during the 1 ^st^ minute in the condition iC1/3s-A against 2^nd^ and 3^rd^ minutes performing the task.

### 4.5. Discussion

The results of this experiment indicate that the introduction of extra beeps caused an increase in cognitive demand for the participants in the intermittent tasks. That is, continuous counting tasks were less demanding than tasks with an intermediate beep. It is possible to conjecture that the intermediate beeps would allow the participant to follow the metronome, and make it easier for them to generate the numbers, but the effect was contrary to this expectation. The only exception to this result was in task iC1/3-A, which was no more demanding than the continuous counting tasks. This result may mean that this task, due to the large gaps between counting, and the very low counting rate, allows the participant not to focus attention on the intermediate beep, or even that the slow pace of this task implies a lower cognitive demand on the part of the participants.

The fact that the iC1/3-B task, although having a three-second count, was different from this one could indicate that rate is one of the determining factors for our results (since it has the same number of beeps as the iC1/1s task), but if this were true, both the iC1/3-B and iC1/1s tasks would not have the same cognitive demand as the iC1/2s task, which has fewer beeps. This result seems to indicate another possible interpretation, that intermediate beeps demand cognitive resources for counting inhibition.

In this experiment, we tried to verify if the manipulation of the automatic tasks transformed them into tasks closer to a controlled demand, also investigating the possibility of a continuum of mental effort (Bugg et al. 2008; Bugg and Crump 2012; Diede and Bugg 2017; Jacoby et al. 2003). The results of the present study can be interpreted as demonstrating that cognitive processing does not follow an explicit dual logic, with some manipulation making easy tasks more demanding, but still not enough for an intense mental effort allocation as discussed in previous studies (Diede and Bugg 2017; Foroughi et al. 2017; Hommel 2007; Konishi et al. 2015).

Both pupil dilation peaks and blinking results corroborate the results found in the first experiment. With regard to peaks, the fact that all tasks are normal, direct, and without a different logic for their accomplishment makes the participants quickly adapt to their achievement, without any surprise or extremely high inhibitory control at the beginning or in the middle of the tasks. With the only exception occurring in the counting conditions of a number per second (with or without intermediate beeps) with an effect on pupil peak. This result demonstrates that continuous counting at a number per second is such a simple task for individuals that apparently when this logic is altered, the attention demand is higher than usual.

The effect of cognitive tasks on blink rates is not apparent in the tasks presented in this study imply that although the use of cognitive resources in the tasks were different, the engagement to perform them was not (Cools and D’Esposito 2011; Wong & Epps, 2016). That is, no great mental effort demand to balance these tasks is apparent. As pointed out in previous studies (Colzato et al. 2008, 2009; Jongkees and Colzato 2016; Konishi et al. 2015), EBR can be related to mind wandering and attentional gaps. In our experiment the participants were concentrated in the accomplishment of the tasks, this may have been a factor for the no-significant difference EBR.

## 5. Conclusion

Controlled and automated tasks demand different mental effort, and this effort is portrayed in pupil dilation. In addition, a simple variable like an intermediate beep can produce a need for inhibition response from the participants, but only when the tasks are faster. These results suggest the possibility of measuring the mental effort of individuals using pupillometric data, allowing differentiation between cognitive systems through the manipulation of automatic tasks.

The results of the second experiment showed that the additional intermediate beep made the counting tasks more demanding. Small manipulations like this can increase the need for inhibitory control.

Intermediate beeps, that initially can be thought as helpers to the participants in the monitoring and implementation of the counting tasks are in fact a distractor, requiring the participants to carry out more inhibitory processing to accomplish these tasks. That is, extra mental effort is needed in performing the tasks

## References

Alnæs D, Sneve MH, Espeseth T, Endestad T, Van de Pavert SHP, Laeng B (2014) Pupil size signals mental effort deployed during multiple object tracking and predicts brain activity in the dorsal attention network and the locus coeruleus. J of Vision, 14(4):1–20. http://www.journalofvision.org/contents/14/4/1. Doi:10.1167/14.4.1

Aston-Jones G, Cohen JD (2005) An Integrative Theory of Locus Coeruleus-Norepinephrine Function: Adaptive Gain and Optimal Performance. Annu Rev Neurosci, 28:403–450. doi:10.1146./annurev.neuro.28.061604.135709

Beatty J (1982) Task-Evoked Pupillary Responses, Processing Load, and the Structure of Processing Resources. Psychological Bulletin, 91(2):276–292.

Beatty J, Lucero-Wagoner B (2000) The Pupillary System. In: John T. Cacioppo, Louis G. Tassinary, & Gary G. Bernston (Eds.), Handbook of Psychophysiology (2 ed.), USA: Cambridge University Press. 142–161.

Berridge CW, Waterhouse BD (2003) The locus coeruleus–noradrenergic system: modulation of behavioral state and state-dependent cognitive processes. Brain Research Reviews, 42:33–84.

Bourisly AK (2015) Pupil Response Diameter is Modulated as a Function of Cognitive Load during Mental Addition: A Psychophysiological Protocol and Study. J of Advance Neuroscience Research, 2:1–6.

Bugg JM, Jacoby LL, Toth JP (2008) Multiple levels of control in the Stroop task. Mem Cognit, 36(8): 1484–1494. doi:10.3758/MC.36.8.1484.

Bugg JM, Crump MJC (2017) In support of a distinction between voluntary and stimulus-driven control: a review of the literature on proportion congruent effects. Front Psychology, 3:367. doi:10.3389/fpsyg.2012.00367

Cabestrero R, Crespo A, Quirós P (2009) Pupillary Dilation as an Index of Task Demands. Perceptual and Motor Skills, 109(3):664–678. doi:10.2466/PMS.109.3.664-678.

Cavanagh JF, Wiecki TV, Kochar A, Frank MJ (2014) Eye Tracking and Pupillometry Are Indicators of Dissociable Latent Decision Processes. Journal of Experimental Psychology: General, 143(4), 1476–1488. doi:10.1037/a0035813.

Cohen J (1994) The earth is round (p < .05). American Psychologist, 49:997–1003. doi:10.1037/0003-066X.49.12.997.

Colzato LS, Slagterb HA, Spapéa MMA, Hommel B (2008) Blinks of the eye predict blinks of the mind. Neuropsychologia, 46:3179–3183. doi:10.1016/j.neuropsychologia.2008.07.006.

Colzato LS, Van den Wildenberg WPM, Van Wouwe NC, Pannebakker MM, Hommel B (2009) Dopamine and inhibitory action control: evidence from spontaneous eye blink rates. Exp Brain Res, 196:467–474.

Cools R, D’Esposito M (2011) Inverted-U-shaped dopamine actions on human working memory and cognitive control. Biol Psychiatry, 69(12):113–125. doi:10.1016/j.biopsych.2011.03.028

Corbetta M, Shulman GL (2002) Control of Goal-Directed and Stimulus-Driven Attention in the Brain. Nature Neuroscience, 3:201–2015. doi:10.1038/nrn755

Christie ST, Schrater P (2015) Cognitive cost as dynamic allocation of energetic resources. Front Neuroscience, 9:289. doi:10.3389/fnins.2015.00289.

Debue N, Van de Leemput C (2014) What does germane load mean? An empirical contribution to the cognitive load theory. Frontiers in Psychology, 5:1–12. doi:10.3389/fpsyg.2014.01099.

Dirnberger G, Jahanshashi M (2010) Response selection in dual task paradigms: observations from random generation tasks. Exp. Brain. Rev, 201:535–548. doi: 10.1007/s00221-009-2068-y.

Diede NT, Bugg JM (2017) Mental effort Is Modulated Outside of the Explicit Awareness of Conflict Frequency: Evidence From Pupillometry. Journal of Experimental Psychology: Learning, Memory, and Cognition. 43(5): 824–835. http://dx.doi.org/10.1037/xlm0000349

Dunlop WP, Cortina JM, Vaslow JB, Burke MJ (1996) Meta-analysis of experiments with matched groups or repeated measures designs. Psychological Methods. 1: 170–177.

Eckstein MK, Guerra-Carrillo B, Singley ATM, Bunge SA (2016) Beyond eye gaze: What else can eyetracking reveal about cognition and cognitive development?. Dev Cogn Neurosci. 25:69–91. http://dx.doi.org/10.1016/j.dcn.2016.11.001

Egner T (2008) Multiple conflict-driven control mechanisms in the human brain. Trends Cogn Sci. 12(10):374–380. doi: 10.1016/j.tics.2008.07.001

Evans FJ (1974) Monitoring attention deployment by random number generation: An index to measure subjective randomness. Bulletin of the Psychonomic Society, 12 (1): 35 – 38.

Foroughi CK, Sibley C, Coyne JT (2017) Pupil size as a measure of within-task learning. Psychophysiology, 54(10):1436–1443. doi: 10.1111/psyp.12896

Fukuda K (1994) Analysis of Eyeblink Activity During Discriminative Tasks. Perceptual and Motor Skills. 79: 1599–1608. http://dx.doi.org/10.2466/pms.1994.79.3f.1599.

Fukuda K, Stern JA, Brown TB, Russo MB (2005) Cognition, Blinks, Eye-Movements, and Pupillary Movements During Performance of a Running Memory Task. Aviation Space and Environmental Medicine, 76: C75–85

Gergelyfi M, Jacob B, Olivier E, Zénon A (2015) Dissociation between mental fatigue and motivational state during prolonged mental activity. Front Behav Neurosci, 9:176. doi:10.3389/fnbeh.2015.00176

Gopher D, Donchin E (1986) Chapter 41. Workload-an examination of the concept. In: Handbook of perception and human performance. New York: Wiley. 1:49.

Gratton G (2018) Brain reflections: A circuit-based framework for understanding information processing and cognitive control. Psychophysiology, 55(33):e13038. https://doi.org/10.1111/psyp.13038.

Hamdan AC, Souza JA, Bueno OFA (2004) Performance of University Students on Random Number Generation at Different Rates to Evaluate Executive Functions. Arq Neuropsiquiatr, 62 (1):58 – 60.doi:http://dx.doi.org/10.1590/S0004-282X2004000100010

Hershaw JN, Ettenhofer ML (2018) Insights into cognitive pupillometry: Evaluation of the utility of pupillary metrics for assessing cognitive load in normative and clinical samples .I J Psychophysiology, 134:62–78. https://doi.org/10.1016/j.ijpsycho.2018.10.008

Hasher L, Zacks RT (1984) Automatic processing of fundamental information: The case of frequency of occurrence. American Psychologist, 39(12):1372–1388.

Hess EH, Polt JM (1964) Pupil Size in Relation to Mental Activity during Simple Problem-Solving. Science, 143:1190–1192. doi: 10.1126/science.143.3611.1190.

Hommel B (2007) Consciousness and control: Not identical twins. Journal of Consciousness Studies, 14:155–176.

Ichikawa N, Ohira H (2004) Eyeblink activity as an index of cognitive processing: Temporal distribuction of eyeblinks as an indicator of expectancy in semantic priming. Perceptual and Motor Skills, 98:131–140.

Isabella SL, Urbain CJ, Cheyne A, Cheyne D (2019) Pupillary Responses and Reaction Times Index Different Cognitive Processes in a Combined Go/Switch Incidental Learning Task. Neuropsychologia, 127, 48–56. https://doi.org/10.1016/j.neuropsychologia.2019.02.007

Jacoby LL, Lindsay DS, Hessels S (2003) Item-specific control of automatic processes: stroop process dissociations. Psychon Bull Rev, 10(3):638–644.

Jainta S, Baccino T (2010) Analyzing the pupil response due to increased cognitive demand: A independent component analysis study. International J of Psychophysiology, 77:1–7. doi: 10.1016/j.ijpsycho.2010.03.008.

Jahanshahi M, Saleem T, Ho AK, Dirnberger G, Fuller R (2006) Random number generation as an index of controlled processing. Neuropsychology, 20:391–399. doi: 10.1037/0894-4105.20.4.391.

Jepma M, Nieuwenhuis S (2011) Pupil diameter predicts changes in the exploration-exploitation trade-off: evidence for the adaptive gain theory. J Cogn Neurosci. 23(7):1587–1596. doi:10.1162/jocn.2010.21548

Jongkees BJ, Colzato LS (2016) Spontaneous eye blink rate as predictor of dopamine-related cognitive function-A review. Neurosci Biobehav Rev. 71:58–82. doi: 10.1016/j.neubiorev.2016.08.020.

Joshi S, Li Y, Kalwani RM, Gold JI (2016) Relationships between Pupil Diameter and Neuronal Activity in the Locus Coeruleus, Colliculi, and Cingulate Cortex. Neuron. 89(1):221–234.

Just MA, Carpenter PA, Miyake A (2003) Neuroindices of cognitive workload: neuroimaging, pupillometric and event-related potential studies of brain work. Theor. Issues in Ergon. Sci. 4 (1-2): 56 – 88. doi: 10.1080/14639220210159735.

Kahneman D (1973) Attention and Effort. Englewoods Cliff, New Jersey: Prentice-Hall.

Kahneman D, Peavler WS (1969) Incentive Effects and Pupillary Changes in Association Learning. Journal of Experimental Psychology. 7 (2): 312–318.

Kang O, Wheatley T (2015) Pupil dilation patterns reflect the contents of consciousness. Consciousness and Cognition, 35:128–135. http://dx.doi.org/10.1016/j.concog.2015.05.001

Kihara K, Takeuchi T, Yoshimoto S, Kondo HM, Kawahara J.I (2015) Pupillometric evidence for the locus coeruleus-noradrenaline system facilitating attentional processing of action-triggered visual stimuli. Front. Psychol. 6:827. doi: 10.3389/fpsyg.2015.00827.

Konishi M, Brown K, Battaglini L, Smallwood J (2015) When attention wanders: Pupillometric signatures of fluctuations in external attention. Cognition. 168: 16–26.

Kool W, Botvinick M (2018) Mental Labour. Nature Human Behaviour. 2: 899–908. https://doi.org/10.1038/s41562-018-0401-9

Laeng B, Sirois S, Gredebäck G (2014) Pupillometry: A Window to the Preconscious. Perspectives on Psychological Science. 7(1):18–27.doi: 10.1177/1745691611427305.

Lee CR, Margolis DJ (2016) Pupil Dynamics Reflect Behavioral Choice and Learning in a Go/NoGo Tactile Decision-Making Task in Mice. Front. Behav. Neurosci. 10:200. doi:10.3389/fnbeh.2016.00200

Leppink J, Pérez-Fuster P (2019) Mental Effort, Workload, Time on Task, and Certainty: Beyond Linear Models. Educational Psychology Review, 31(2):421–438. https://doi.org/10.1007/s10648-018-09460-2

Massar SAA, Lim J, Sasmita K, Chee MWL (2016). Rewards boost sustained attention through higher effort: A value-based decision making approach. Biological Psychology. 120:21–27. http://doi.org/10.1016/j.biopsycho.2016.07.019.

Mathot S, Fabius J, van Heus E, Van der Stigchel S (2018) Safe and sensible preprocessing and baseline correction of pupil-size data Behavior Research Methods. 50(1):94–106. https://doi.org/10.3758/s13428-017-1007-2.

McCabe DP, Roediger HL, Karpicke JD (2011) Automatic Processing Influences free recall: conveging evidence from the process dissociation procedure and remember-know judgments. Mem. Cogn. 39:389–402. doi 10.3758/s1321-010-0040-5.

Miyake A, Friedman NP, Emerson MJ, Witzki AH, Howerter A, Wager TD (2000) The unity and diversity of executive functions and their contributions to complex “frontal lobe” tasks: a latent variable analysis. Cogn. Psychol. 41:49–100.

Mitra R, McNeal KS, Bondell HD (2017) Pupillary response to complex interdependent tasks: A cognitive-load theory perspective. Behav Res, 49:1905–1919. doi:10.3758/s13428-016-0833-y

Moresi S, Adam JJ, Rijcken J, Van Gerven PWM, Kuipers H, Jolles J (2008) Pupil dilation in response preparation. I J Psychophysiology, 67:124–130.

Murphy PR, O’Conneel RG, O’Sullivan M, Robertson IH, Balster JH (2014) Pupil diameter covaries with BOLD activity in human brain locus coeruleus. Human Brain Mapping. 35(8):4140–4154.

Neuringer A (1986). Can People Behave “Randomly?”: The Role of Feedback. J Exp Psy Gen. 115 (1), 62 – 75. http://dx.doi.org/10.1037/0096-3445.115.1.62

Oliva M (2019) Pupil size and search performance in low and high perceptual load. Cognitive, Affective & Behavioral Neuroscience 19:366–376. https://doi.org/10.3758/s13415-018-00677-w

Paas F, Ayres P (2014) Cognitive Load Theory: A Broader View on the Role of Memory in Learning and Education. Educ Psychol Rev. 26:191–195. doi: 10.1007/s10648-014-9263-5.

Posner MI, Fan J (2008) Attention as an organ system. In: Pomerantz JR, editor. Topics in Integrative Neuroscience: From Cells to Cognition. Cambridge University Press. 31–61.

Querino M, Santos L, Ginani G, Nicolau E, Miranda D, Romano-Silva M, Malloy-Diniz L (2015) Mental effort and pupil dilation in Controlled and automatic processes. Translational Neuroscience. 168–173. doi:10.1515/tnsci-2015-0017.

Rondeel EWM, Van Steenbergen H, Holland RW, Van Knippenberg A (2015) A closer look at cognitive control: differences in resource allocation during updating, inhibition and switching as revealed by pupillometry. Front. Hum. Neurosci. 9:494. doi:10.3389/fnhum.2015.00494.

Schneider W, Shiffrin RM (1977) Controlled and Automatic Human Information Processing: I. Detection, Search, and Attention. Psychological Review. 84 (1):1–66.

Schulz MA, Schmalbach B, Brugger P, Witt K (2012) Analyzing Humanly Generated Random Number Sequences: A Pattern-Based Approach. PLOS ONE. 7(7): E41531.doi:10.1371/jornal.pone.0041531.

Shiffrin RM, Schneider W (1977) Controlled and Automatic Human Information Processing: II. Perceptual Learning, Automatic Attending, and a General Theory. Psychological Review. 84(2):127–190.

Siegle GJ, Ichikawa N, Steinhauer S (2008) Blink before and after you think: Blinks occur prior to and following cognitive load indexed by pupillary responses. Psychophysiology.45, 679–687.doi:10.1111/j.1469-8986.2008.00681.x.

Sirois S, Brisson J (2014) Pupillometry. WIREs Cogn Sci.doi:10.1002/wes.1323.

Steiger JH (2004) Beyond the F Test: Effect Size Confidence Intervals and Tests of Close Fit in the Analysis of Variance and Contrast Analysis. Psychological Methods. 9 (2):164–182.

Sweller J (1988) Cognitive Load During Problem Solving: Effects on Learning Cognitive Science. 12:257–285. doi: 10.1207/s15516709cog1202_4.

Towse JN, Neil D (1998) Analyzing human random generation behavior: A review of methods used and a computer program for describing performance. Behavior Research Methods, Instruments, & Computers. 30 (4):583–591.

Van Bochove M, Van der Haegen L, Notebaert W, Verguts T (2013) Blinking predicts enhanced cognitive control. Cogn Affect Behav Neurosci. 13:346–354.doi: 10.3758/s13415-012-0138-2.

Van Der Mer E, Beyer R, Horn J, Foth M (2010) Bornemann B, Ries J, Kramer J, Warmuth B, Heekeren HK, & Wartenburger I. Resource allocation and fluid intelligence: Insights from pupillometry. Psychophysiology. 47:158–169. doi: 10.1111/j.1469-8986.2009.00884.x.

Van der Wel P, van Steenbergen H (2018) Pupil dilation as an index of effort in cognitive control tasks: A review. Psychonomic Bulletin & Review 25:2005–2015. https://doi.org/10.3758/s13423-018-1432-y

Wierda SM, Van Rijn H, Taatgen NA, Martens S (2012) Pupil dilation deconvolution reveals the dynamics of attention at high temporal resolution. PNAS. 109: 8456–8460.

Wong HK, Epps J (2016) Pupillary transient responses to within-task cognitive load variation. 137:47–63. doi: 10.1016/j.cmpb.2016.08.017

Yeo G, Neal A (2008) Subjective Cognitive Effort: A Model of States, Traits, and Time. Journal of Apllied Psychology. 93(3): 617–631. doi:10.1037/0021-9010.93.3.617

Zhang T, Mou D, Wang C, Tan F, Jiang Y, Lijun Z, Li H (2015) Dopamine and executive function: increased spontaneous eye blink rates correlate with better set-shifting and inhibition, but poorer updating. Int J Psychophysiol, 96:155–161. doi: 10.1016/j.ijpsycho.2015.04.010

